# Co-evolutionary dynamics of mammalian brain and body size

**DOI:** 10.1101/2023.06.08.544206

**Authors:** Chris Venditti, Joanna Baker, Robert A. Barton

## Abstract

Despite decades of comparative studies, fundamental questions remain about how brain and body size co-evolved. Divergent explanations exist concurrently in the literature for phenomena such as variation in brain relative to body size, variability in the scaling relationship across taxonomic levels and between taxonomic groups, and purported evolutionary trends. Here we resolve these issues using a comprehensive dataset of brain and body masses across the mammal radiation, and a method enabling us to study brain relative to body mass evolution while estimating their evolutionary rates along the branches of a phylogeny. Contrary to the rarely questioned assumption of a log-linear relationship, we find that a curvilinear model best describes the evolutionary relationship between log brain mass and log body mass. This model greatly simplifies our understanding of mammalian brain-body co-evolution: it can simultaneously explain both the much-discussed taxon-level effect and variation in slopes and intercepts previously attributed to complex scaling patterns. We also document substantial variation in rates of relative brain mass evolution, with bursts of change scattered through the tree. General trends for increasing relative brain size over time are found in only three mammalian orders, with the most extreme case being primates, setting the stage for the uniquely rapid directional increase that ultimately produced the computational powers of the human brain.

---

Here we use a phylogenetic approach applied to a comprehensive dataset of brain and body masses (n=1504) spanning the mammalian radiation to flexibly characterize the underlying brain-body mass (BBM) relationship whilst simultaneously detecting rapid bursts or reductions in the rate of relative brain mass evolution^1-3^ (Methods). The BBM relationship across diverse animal groups, such as the mammals, is often studied by fitting models with multiple slopes and intercepts to account for differences between clades ^e.g. 4,5-7^. In congruence with previous studies, we find that a model which allows the BBM relationship to vary within taxonomic groups (multiple-slopes) fits our data better than a single BBM relationship (single-slope) in a log-log space (Bayes factor [*BF*] = 161.198, Figure 1).

**Figure 1:**
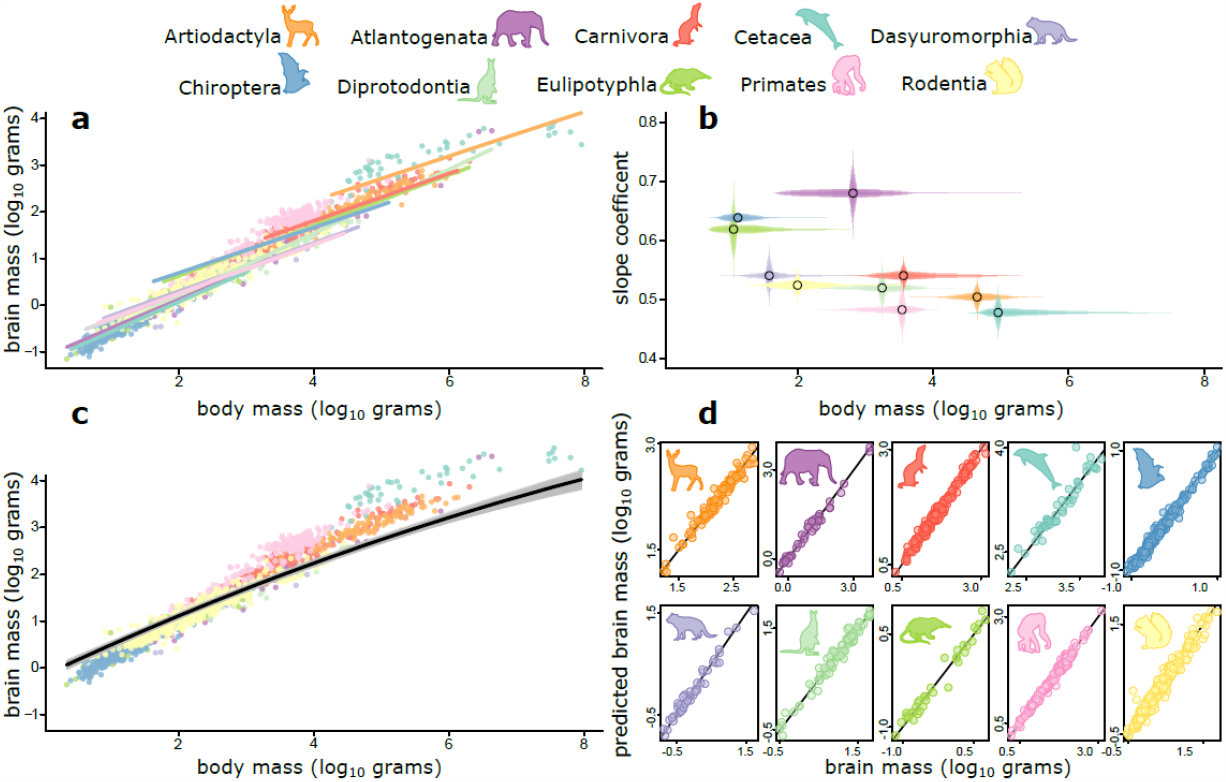
The curvilinear relationship of the brain-body mass (BBM) relationship across mammals. a) The BBM relationship across mammals with median model predictions from the variable rates regression model including separate intercepts and slopes for taxonomic groups. b) slope coefficient (and percentiles of posterior distributions plotted as transparent lines) for each taxonomic group from panel a, plotted against body mass (percentiles of body mass range plotted as transparent lines) for each group. c) The BBM relationship across mammals with model predictions from the variable rates regression model including the best fit curvilinear relationship (the posterior distribution of curves is shown in grey, median in black). d) Actual brain mass against predicted brain mass from the curvilinear model highlighting the accuracy of the fit to the data. In all panels, points and lines are coloured in accordance with orders as shown by the representative silhouettes (see legend, not to scale).

However, across the range of mammalian body mass, there is a negative correlation between the slope of a mammalian taxonomic group and its average body mass (ρ=-0.63, p=0.049, figure 1b), indicating potential mass-dependency for the BBM relationship (ρ=-0.82, p=0.006 after excluding Atlantogenata which spans the extremes of mammalian body size range - over 5 orders of magnitude). There have been some hints in the literature over the years which point to a potential mass-dependency for the BBM relationship^8-10^. However, this is largely ignored by researchers and has never been tested against a multiple slopes model nor in a phylogenetic context. To explicitly test between alternatives, we fit a curvilinear relationship (second order polynomial) to test for this mass-dependency. This curvilinear mass-dependent model fits significantly better than the multiple-slope model (*BF* = 264.461, Figure 1c): as mammals increase in mass, the rate at which brain mass increases with body mass decreases (this result is robust to the results are robust to intraspecific variation). There is no significant variation in the curvilinear relationship amongst taxonomic groups and the single quadratic slope fits the data extremely well across all mammalian groups with no systematic bias (Figure 1d) . Thus, phenomena such as the previously reported variability in the slopes and intercepts of the log-linear BBM relationship across mammalian orders ^e.g.7^ and evolutionary lags in brain mass relative to body mass^e.g.11^ are explained exclusively as mass-dependent effects rather than taxon-specific patterns of brain evolution.

With this mass-dependence in mind, we can also shed new light on the well-known ‘taxon-level effect’ ^e.g.12^ in the BBM relationship among mammals. The taxon-level effect is an enduring idea, describing a phenomenon where the slope of the BBM relationship is shallower when investigated across closely related species, such as those in the same genus, compared to across distantly related species (such as families or orders). Although many ideas to explain the taxon-level effect have been proposed^10,12-15^, none have proven robust, and for that reason the phenomenon and its causes remain contentious. However, we suggest that the apparent taxon-level effect emerges simply as a side-effect of the curvilinearity of the BBM relationship combined with the trend for increase in body mass over time, known as Cope’s rule^16^ . Strong evidence from both the fossil record^17^ and extant species^1,18^ support Cope’s rule. Given this pattern, linear regression coefficients will inevitably be shallower in more closely related species compared to more distantly related species. This is purely because more distantly related species are more likely to have branches that span into deep time such that the evolutionary signature of Cope’s rule should be stronger. Our results therefore suggest that more complex evolutionary explanations for the taxon-level effect, involving decoupling of brain and body mass and evolutionary lags^e.g.10,13^, are unnecessary.

Even after accounting for the mass-dependent scaling of the BBM relationship, there is substantial variation in evolutionary rates – the rate of relative brain mass evolution is highly heterogeneous (Figure 2). All mammalian groups show branches where the rate of relative brain mass evolution is increased; this is most pronounced in Primates, Rodentia, and Carnivora (Figure 2), contrary to suggestions that relative brain mass predominantly reflects body mass evolution (e.g.^7,10,19,20^).

**Figure 2:**
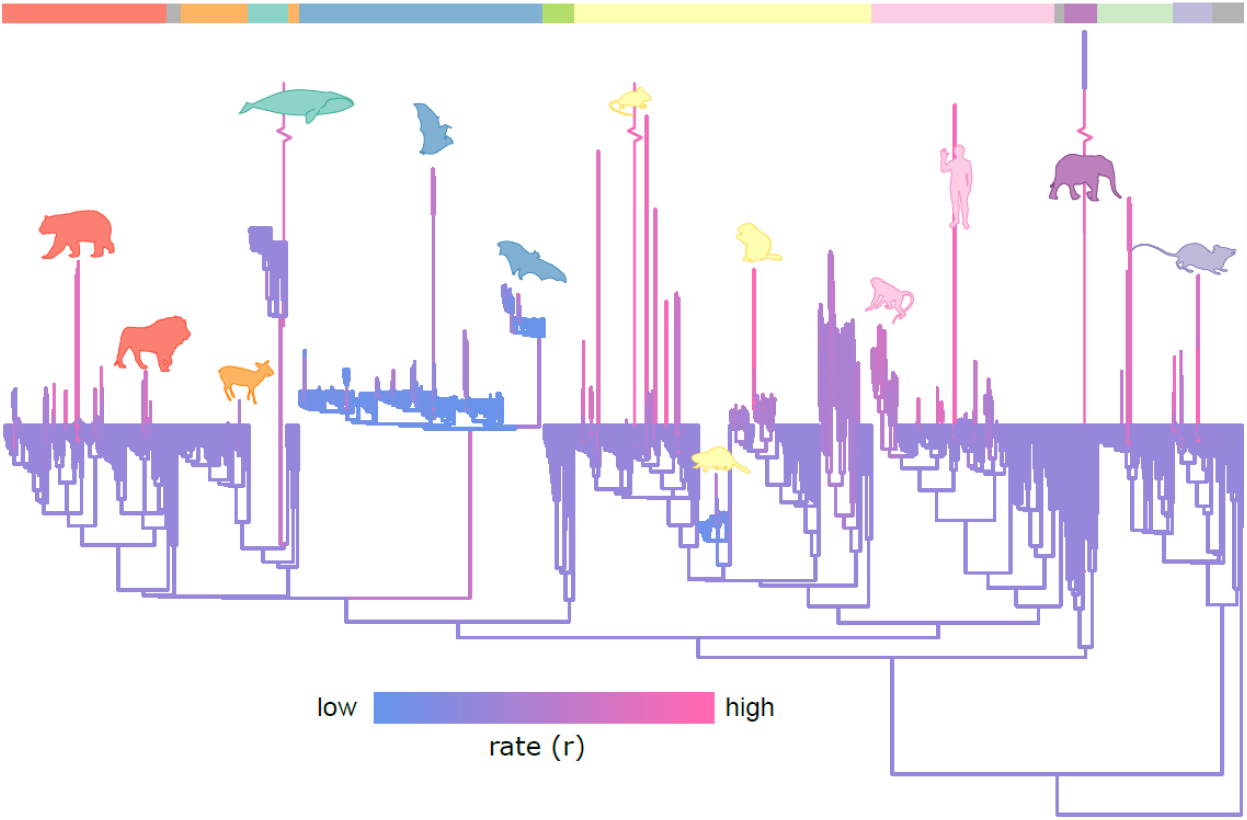
Rates of relative brain mass evolution. The mammal phylogenetic tree used in this study where branches are coloured, compressed, and stretched by the median relative rate of brain mass evolution. Mammalian taxonomic groups are represented by coloured bars along the top of the figure which correspond to the colours of the silhouettes in Figure 1. Selected branches are highlighted by representative silhouettes using the same colour scheme (not to scale).

Although there is a high rate on the branch leading to Chiroptera (Figure 2), bats as a clade tend to have a very low rate of relative brain mass evolution (∼2.5 times lower than the mammalian background rate), which might indicate an evolutionary constraint associated with flight. Within bats, clades that show significantly elevated rates are not united by any obvious ecological factor such as diet – which has previously been suggested to have been a driver of brain mass evolution in bats^21^. Hence, more work is needed to understand the drivers of these variable rates. Confirming previous suggestions^22^, the rate of relative brain mass change we observe on the branch leading to humans is extremely high, with a median rate 22.9 times higher than the mammalian background rate.

Analyses of rate heterogeneity like those we use here introduce meaningful variation into the branch lengths of a phylogeny (Figure 2). This makes it possible to study evolutionary trends in trait (or relative trait) evolution through time ^e.g.1,23,24^. Longer branches represent an increase in the rate of evolution likely owing to the influence of selection^24,25^ - they have undergone more relative brain mass change than would be expected given their length in time. The sum of all rate-scaled branches along the evolutionary path to each species (root-to-tip rate) can therefore be used to measure the total amount of adaptive change that a species has experienced during its history^1,24^. With this in mind, we use Bayesian phylogenetic regression models to determine if there have been any long-term evolutionary trends in relative mammalian brain mass through time (e.g., towards larger or smaller mass). Across all mammals we find a significant increase in relative brain mass with root-to-tip rate (β=0.906, *P*_*x*_=0.000). However, when we allow the relationship to vary among orders, we find a significant trend in only three orders: rodents (β=0.998, *P*_*x*_=0.002), carnivores (β=1.845, *P*_*x*_=0.000) and, most strikingly, primates (β=2.074, *P*_*x*_=0.000), (Figure 3). No such relationship is found in any other order or across all remaining mammals (*P*_*x*_= 0.331). This demonstrates that the Marsh-Lartet rule^26,27^, proposing a trend for relative brain mass to increase throughout mammalian evolution, is not a general mammalian phenomenon seen in all orders.

**Figure 3:**
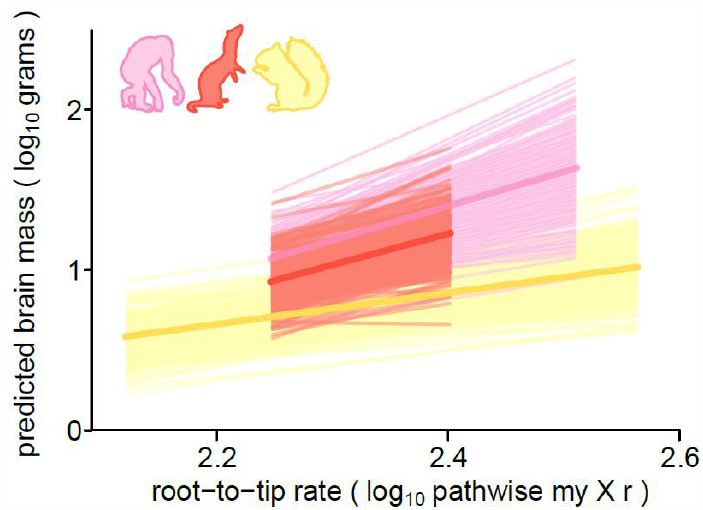
Trends towards increasing brain mass through time. Posterior distributions (transparent lines) and median (solid line) of the model predictions demonstrate the trend in three mammalian orders (rodents, yellow; carnivores, red; primates, pink). Silhouettes indicate the relevant taxonomic groups (see Figure 1) and are not to scale.

The unique trajectory we reveal in primates is apparent if we simply compare the inferred body mass and brain mass change along the branches of the phylogenetic tree for each taxonomic group of mammals we study (Figure 4, estimated using the parameters of our models; see methods). When the proportion of branches where we observe brain mass increase are compared with the proportion where body mass increases, only the primates and carnivores are a clear positive outlier. Primates are the extreme case in which over 80% of branches underwent a brain mass increase, compared with under 65% where body mass increased (figure 4). We can further examine the unique nature of this trend when we compare the standardised magnitude of change in brain compared to body. We conducted an analysis of covariance accounting for ancestry, comparing the reconstructed brain change along each branch in each of our mammalian groups, accounting for body size change (p<0.0001, figure 4). The inset of figure 5 shows the result of a post hoc Tukey HSD (honestly significant difference) test, which demonstrates (along with the coloured radial trees in Figure 5) that among mammals, primates have the highest relative change in brain size followed by carnivores and rodents. Because the trend in relative size is clearly not driven exclusively by body size change, we can therefore exclude the hypothesis that large relative brain size in primates predominantly reflects reductions in body size^10^.

**Figure 4:**
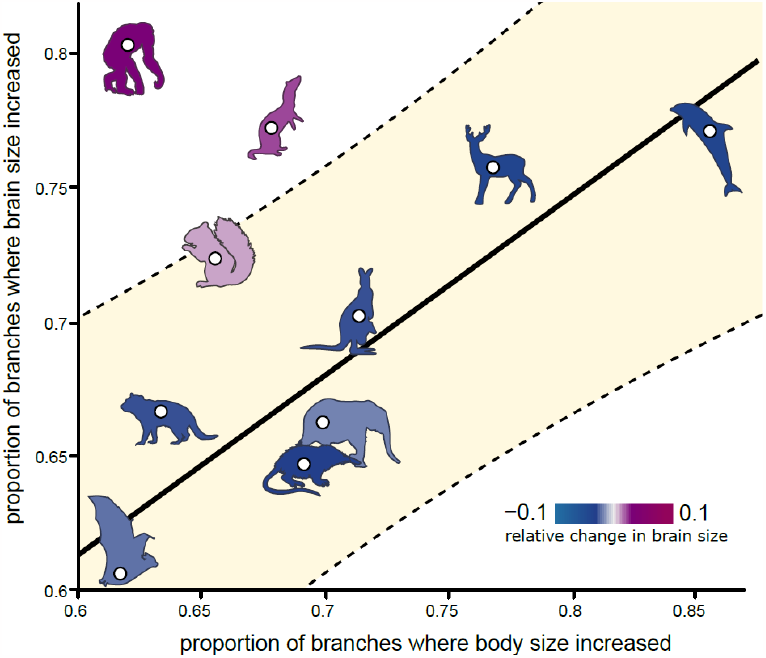
Proportion of branch-wise brain and body size increases. The proportion of branches where brain mass increased compared with the proportion of branches where body mass increased is plotted per taxonomic group. The fitted line and 95% prediction intervals (shaded) highlight primates and carnivores are outliers. A representative silhouette for each taxonomic group (see Figure 1) is shown and coloured according to the total relative change in brain size (estimated difference in the standardised magnitude of change in brain mass compared to body mass, *Z*_*brain*_*-Z*_*body*_).

**Figure 5:**
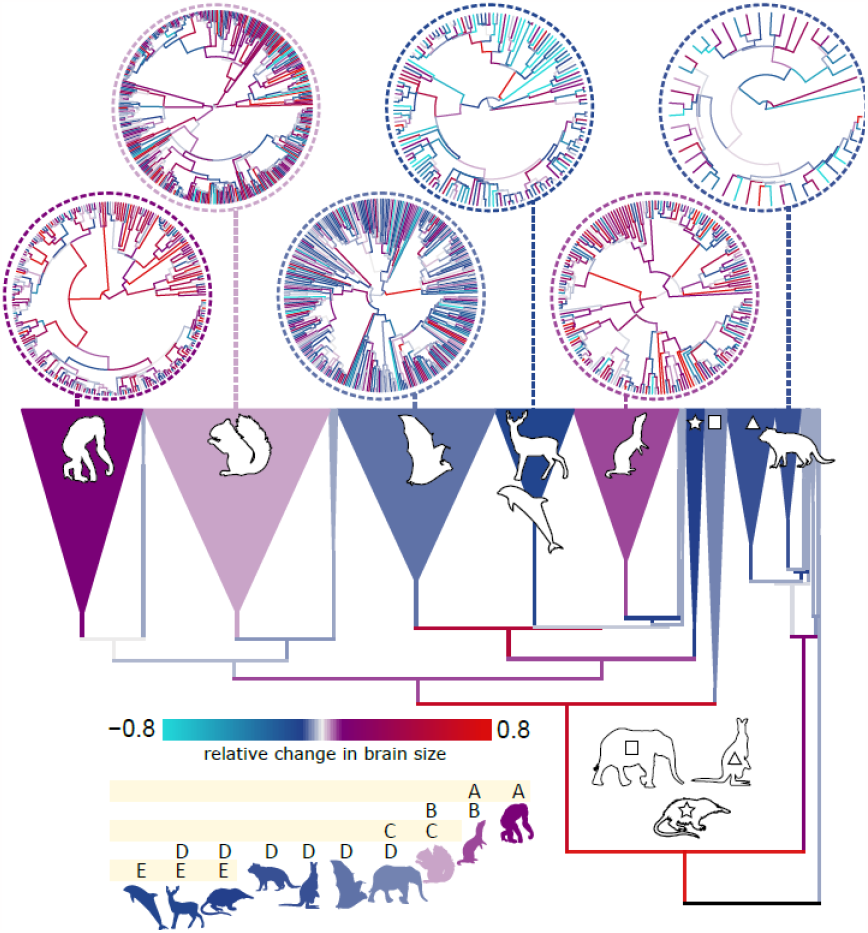
Magnitude of branch-wise brain and body change. Phylogenetic branches (and collapsed clades) are coloured by the estimated difference in the standardised magnitude of change in brain mass compared to body mass (Z_brain_-Z_body_). Inset shows the result of a post hoc Tukey HSD (honestly significant difference) test. Silhouettes for each of the clades we study (see Figure 1) are displayed on the collapsed portion of the phylogeny (symbols are shown for Atlantogenata [square], Diprotodontia [triangle] and Eulipotyphla [star] to facilitate clear visualization of the phylogeny).

Natural selection has decoupled brain and body mass evolution in primates to a unique extent, producing sustained and directional increase in relative brain mass for over 55 million years. This trend set the stage for rapid increase in hominins, leading ultimately to modern humans’ unprecedentedly large brains^22,28^. Hence, the emergence of human-like cognitive capacities was facilitated by a shift in the fundamental pattern of brain evolution at the origin of the primates. An explanation for this distinctive pattern in primates – whether in terms of the release of a constraint or an adaptive shift that initiated an escalating feedback between brain and behaviour - would be a significant contribution to biology. Candidates might include sociality^29^, unusual patterns of maternal investment facilitating extended brain growth^30^, the advent of visuo-motor control of the forelimb associated with stereoscopically guided grasping and manipulation and a unique pattern of connectivity between eye and brain^31-33^.

Our results have significant implications for the study of brain size evolution. By simultaneously explaining multiple statistical phenomena that had been reported on the basis of linear models, our results resolve several debates about the co-evolutionary dynamics of mammalian brain and body size. This obviates the need for special explanations proposed for each individual phenomenon, including the taxon-level effect, lag in brain mass relative to body mass evolution, differences in brain-body intercepts (‘grade shifts’) and differences in slopes^6,7,34,35^. Previous conclusions regarding the evolutionary relationship between brain and body mass, how it changes through time and or among phylogenetic groups, estimates of relative brain mass in particular taxa and methods for deriving them e.g.^7,20,27^ will need to be re-evaluated in light of our findings.

Our results can explain why estimates of encephalization are biased with respect to mass, being lower than expected in large-bodied mammals^27^. In addition, our results suggest that correlates of relative brain mass evolution across taxa varying substantially in mass – whether behaviour, ecology development or life history - will need to be re-assessed after accounting for the curvilinearity in the BBM relationship. The evolutionary routes to large relative brain size in extant species were variable, with no single process applying across the mammalian tree. The questions of why a curvilinear relationship exists (e.g. whether this reflects the way in which neuronal parameters or connectivity scale with size), and how these findings may apply to other taxonomic groups, will be interesting ones for future investigation.

## Methods

### Data and phylogenetic tree

Our primary data set comprised of 1504 brain and body masses taken from the literature (Table S1). To check our principal results were robust to intraspecific variation we re-ran the analyses using the dataset of Tsuboi *et al*^15^ (n_species_=919, n_samples_=1908) which samples multiple (between 1 and 59) brain and body size data per species using the package MCMCglmm^36^, the results were quantitatively identical: A curvilinear model is still strongly supported (p < 0.001 for the quadratic parameter, estimated in a gaussian model with alpha-expanded priors on the phylogenetic variance). The phylogenetic tree was taken from the time tree of life (Hedges et al. 2015).

### Rate heterogeneity

To determine the extent of variation in the rate of brain mass evolution after accounting for body mass (relative brain mass evolution) we used the *variable rates regression model* ^2,25^. This Bayesian Markov chain Monte Carlo regression technique acts to estimate the rate of evolution in the phylogenetically structured residual error of a regression model^25^. The model simultaneously estimates a Brownian motion process (background rate,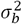) with a set of rate scalars *r* defining branch wise shifts (identifying branches evolving faster (*r >* 1) or slower (0 ≤ *r*<1) than the background rate). We then multiply the original branch lengths (measured in time) by the corresponding *r* for each branch, resulting in a scaled phylogeny where longer branches (compared to their original length in time, *r >* 1) indicate faster rates of morphological evolution, and shorter branches (0 ≤ *r*<1) have slower rates. These branch-specific scalars therefore optimize the fit of the phylogeny to the underlying background rate 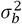 given the inferred phenotypic change along each branch.

To identify evidence for rate heterogeneity we used Bayes Factors (*BF, BF* = −2 log_*e*_[*m*_1_/*m*_0_],), comparing the marginal likelihood of our variable rates model (*m*_1_) to that of a model with a single underlying 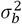 (*m*0). We estimated marginal likelihoods using stepping-stone sampling ^37^ in BayesTraits^38^. Ran 200 stones with 1 million iterations drawing values from a beta-distribution (α= 0.40, β= 1)^37^ and discarded the first 250,000 iterations to ensure convergence. The variable rates model is implemented within a Markov Chain Monte Carlo framework, giving us a posterior distribution of estimated *r* and 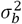. Results were replicated over multiple independent chains.

### Characterising the brain-body mass relationship

We determined the best fitting BBM relationship using Bayes Factors as described above. We identified significant rate heterogeneity in all models (single slope, multiple slopes, and the curvilinear model). Following previous studies, our multiple slopes model was constructed to fit a separate slope and intercept for each mammalian order. However, we only do this where we have more than 20 representatives of that order in our data owing to the suggestion that one should have at least 10 data points per parameter estimated^39^ (we estimate a slope and intercept per group thus we need N>=20). We calculate the proportion of the posterior distribution of each regression parameter that crosses zero (*P*_*x*_). Where *P*_*x*_< 0.05, this means that less than five percent of the posterior distribution crosses zero, and we consider a variable to be substantially different from zero.

### Directionality in relative brain mass evolution

Our method of detecting rate heterogeneity makes it possible to study evolutionary trends in trait evolution^40^ owing to the fact it introduces biologically meaningful variation into the branch lengths of a phylogeny. Longer branches indicate an increase in the rate of evolution most likely arising from selective forces^25,41^; they have experienced more relative brain mass change than would be expected given their length in time. The sum of all rate-scaled branches along the evolutionary path of a species (*path-wise rates*) can therefore be used to measure the total amount of adaptive change that species has experienced during its history ^40^. We used this logic to determine whether there have been any long-term evolutionary trends in relative brain mass evolution and whether they differ among mammalian orders.

We performed all trends analyses using Bayesian phylogenetic regression. We used the median path-wise rate as our predictor variable (but results do not qualitatively differ using the mean or mode). We assessed significance of parameters using the *P*_*x*_< 0.05 criterion described above. All trends analyses were conducted on the median rate-scaled phylogeny in order to account for differences in the amount of brain mass change expected owing to rate heterogeneity ^40^.

### Branch-wise magnitudes and proportions of change

In order to estimate the amount of brain and body mass evolution along each branch of the phylogeny, we used a phylogenetic predictive modelling approach as described in Baker et al^1^. This approach allows us to account for not only the relationships we detect here (curvature and trends in brain size) but also rate heterogeneity and a generalized tendency for body mass to increase through time (Cope’s rule)^1^. We first reconstructed body mass at each node of the phylogeny whilst accounting for the known relationship between body mass and the rate of body mass evolution across mammals^1^. To do this, we ran a *variable rates* model^2^ estimating the rate of body mass evolution across the mammal phylogeny (N = 1504). We then imputed body mass at each node of the phylogenetic tree using the inferred maximum-likelihood relationship between body mass and the median-root-to-tip rate from this analysis (β = 0.009, α = 1.07, *p* < 0.001). We then used these body masses to impute ancestral brain mass at each internal node using the median estimated parameters of our BBM plus trends (see^1^ for details). These reconstructed brain and body sizes provide a realized visualisation of our phylogenetic statistical models.

We then tracked rates, body mass change and brain mass change on a branch-by-branch basis across the phylogeny. For each branch, we calculated the magnitude and direction of change for both brain and body mass from start to end. We then estimated the overall proportion of change by dividing ancestral mass by descendant mass accounting for the time elapsed along the branch.

